# TRACEY: an updated resource for SNARE protein domain annotation with improved HMMs and expanded sequence coverage

**DOI:** 10.64898/2026.06.08.729283

**Authors:** Carlos Pulido-Quetglas, Dirk Fasshauer

**Affiliations:** Department of Computational Biology, University of Lausanne, Lausanne, CH-1015, Switzerland

## Abstract

**Motivation:** SNARE proteins catalyse membrane fusion across the eukaryotic endomembrane system, from synaptic vesicle exocytosis to intracellular trafficking, endosomal and vacuolar transport, and autophagy, and their accurate domain annotation depends on the quality of profile models and the sequence diversity behind them. The original SNARE domain classification predates the recent expansion of eukaryotic sequence data, leaving its HMM profiles and subgroup coverage unable to resolve divergent and lineage-specific paralogs.

**Results:** We present an updated release of TRACEY built on a resynchronized, non-redundant collection of 18,915 curated SNARE proteins spanning 1,188 species, together with a consolidated set of 83 HMM profiles, including 43 models for newly defined subgroups, reconstructed through an iterative, mixture-model-driven procedure. In direct comparison with the legacy models, at least ∼75% of sequences in every overlapping group scored better with the new HMMs, indicating systematic gains in domain detection. A redesigned web interface adds multiparameter querying, FASTA download, and direct scanning of user-submitted sequences against the curated profiles.

**Availability and implementation:** TRACEY is freely available at https://tracey.unil.ch.

**Supplementary information:** Supplementary data are available at Bioinformatics online.

## Introduction

The SNARE (Soluble N-ethylmaleimide-sensitive factor Attachment protein REceptor) protein family catalyses membrane fusion throughout the eukaryotic endomembrane system. Beyond the well-known role of neuronal SNAREs in synaptic vesicle exocytosis and neurotransmitter release (Jahn and Fasshauer, 2012), SNARE-mediated fusion underlies essentially every intracellular trafficking step, including ER-to-Golgi and intra-Golgi transport, endosomal sorting, vacuolar and lysosomal delivery, and autophagy (Jahn and Scheller, 2006). This broad involvement makes SNAREs central not only to secretion but to membrane traffic and organelle homeostasis across the cell. TRACEY consolidates information on SNARE protein domains into a single resource, supporting the comparative, functional, and evolutionary studies that depend on accessible, well-organized sequence data.

The original SNARE domain database by **Kloepper *et*□*al***. (2007) provided the first unified, phylogeny-based classification of eukaryotic SNARE proteins into conserved groups and subgroups, building on the structural reclassification of SNAREs into Q- and R-types (Fasshauer et al., 1998), and it has since served as a broadly adopted reference for functional annotation, comparative genomics, and evolutionary analyses. Its curated domain definitions and hierarchical classification framework remain central to studies of membrane-trafficking machinery. Nonetheless, its scope reflects the sequence space available at the time. The substantial expansion of available eukaryotic genomes, together with the recognition of lineage-specific paralogs and divergent subgroup structure (Kienle et al., 2009), now exceeds the coverage and resolution of the original release. These limitations, particularly in the sensitivity of the original HMM profiles and the restricted sequence diversity they were built on, now constrain accurate subgroup assignment and comparative analysis, underscoring the need for an updated resource to provide a modern, comprehensive, and interoperable reference while maintaining continuity with the established nomenclature.

Since the original release, public sequence repositories have expanded dramatically (Sayers et al., 2024). This increase in available data has revealed extensive SNARE diversity beyond the range captured by the original database. Additionally, evolving standards for database usability, annotation practices, and interface design highlight the need for improved search capabilities, more consistent subgroup assignment, and streamlined access. Together, these developments necessitate a comprehensive update that integrates expanded sequence content, improved HMM models, and a refined subgroup classification system to support accurate annotation and robust evolutionary analysis.

Synchronization of the original TRACEY dataset with current NCBI records resulted in the deprecation of 3,633 sequences, replacement of 1,715 sequences with updated versions, and the addition of 911 newly identified SNARE proteins, highlighting the extent to which the original dataset no longer reflects current sequence space.

TRACEY addresses these limitations by substantially expanding the underlying sequence collection. The database now includes refined HMM profiles built through an updated iterative procedure, enabling improved sensitivity for divergent sequences and supporting a revised subgroup structure grounded in HMM-based evidence. In addition, TRACEY introduces new functionality that allows users to query their own sequences directly against the curated HMMs, facilitating reproducible classification and streamlined annotation. Together, these advances not only provide a more comprehensive and discriminative reference resource, but also enable users to scan external sequence repositories to identify SNARE candidates from lineages not yet represented in TRACEY, accelerating the discovery of missing or under sampled taxonomic groups.

### System and methods

TRACEY currently comprises 18,915 curated SNARE protein sequences (the live database entries retained after the NCBI synchronization described below), with 19,947 annotated SNARE domains, spanning 1,188 species across 581 taxonomic families. Taxonomy is synchronized with NCBI via E-utilities (Sayers, 2009) to ensure up-to-date lineage assignment. Each SNARE protein domain is assigned to one of 83 SNARE domain groups, for which we provide optimized HMM profiles. Across the eukaryotic tree, the collection is dominated by Opisthokonta (Metazoa, 498 species; Fungi, 392 species), with Viridiplantae contributing 108 species; the remaining 190 species span a broad range of protist lineages, including the SAR supergroup (83 species), Discoba (31), protist opisthokonts such as choanoflagellates (19), Amoebozoa (18), Metamonada (11), red algae and glaucophytes (Archaeplastida; 7), Haptista (4), Cryptista (1) and Malawimonada (2). A full breakdown of taxonomic coverage by kingdom and eukaryotic supergroup is provided in Supplementary Table S2. The canonical SNARE repertoire is organized into four main groups (Qa, Qb, Qc, R), each partitioned into classes I–IV and further into 52 subclasses; in addition, TRACEY includes special categories outside this scheme, notably R.Reg (R group; 3 subclasses) and SNAP (two classes, b and c, totalling 6 subclasses) (Kloepper et al., 2007). Compared with the previous release, we refined the 40 existing HMMs and introduced 43 new HMMs corresponding to newly defined subclasses, yielding a consolidated set of 83 models.

### Algorithm

#### HMM model optimization

Profile Hidden Markov Models (HMMs) for the different SNARE groups and subgroups were reconstructed using an iterative, data-driven procedure that progressively expands the training sequence set while applying a controlled inclusion threshold at each iteration to limit low-confidence recruitment.

#### Initial phylogenetic tree reconstruction

Phylogenetic trees were reconstructed independently for each major SNARE class (e.g. Qa.I, Qa.II, Qa.III, Qa.IV), based on the current TRACEY classification. Only the protein regions corresponding to the SNARE domain were used for tree inference. All subsequent analyses for HMM optimization were performed exclusively on domain sequences to ensure consistent homology assessment.

Multiple sequence alignments were generated separately for each SNARE class and used for maximum-likelihood tree reconstruction with IQ-TREE (Minh et al., 2020). The best-fitting amino acid substitution model was selected automatically, and branch support was assessed using 1,000 ultrafast bootstrap replicates and 1,000 SH-aLRT tests. Trees were inferred using the following command:

iqtree -s <class>.alignment --seqtype AA -m TEST -bb 1000 -alrt 1000 --prefix <class>

The resulting domain-based phylogenetic trees provided the reference framework for initial sequence selection and iterative HMM refinement.

#### Initial seed selection

For each SNARE group and subgroup, an initial set of seed sequences was selected based on the current classification of TRACEY entries and the corresponding phylogenetic tree previously described. Seed sequences were preferentially chosen from well-studied model organisms, including Homo sapiens, Mus musculus, Saccharomyces cerevisiae, and a limited number of additional taxa with well-annotated SNARE repertoires (Supplementary Fig. S1-S15). This curated initial set was used to ensure high confidence in group assignment during the first iteration.

#### Iterative HMM refinement

Initial seed sequences were first aligned using MUSCLE (Edgar, 2022), with the -super5 option, which is optimized for accurate alignment of large and diverse sequence sets, and used as the first testing set of sequences. A temporary HMM was constructed from this alignment using standard HMMER procedures (Eddy, 2011) to be used in the first iteration.

At each iteration, sequences included in the training set were realigned using MUSCLE. The resulting alignments were used to reconstruct updated HMMs. This model was then used to scan the complete sequence set associated with the corresponding SNARE group tree using hmmsearch. For each iteration, candidate sequences were selected based on their statistical support and added to the testing set. This procedure was repeated iteratively until convergence, defined as the iteration at which no new sequences were recruited above the selection threshold. The main loop was bounded by a hard limit of 11 iterations; when an iteration recruited no additional sequences, the inclusion threshold was progressively relaxed for up to 30 successive attempts before convergence was declared, ensuring that the threshold space was adequately explored and preventing premature stopping due to numerical fluctuations of the GMM fit. In practice, most groups converged within three iterations, two groups required six and seven, and three groups (Qc.III.c, Qc.III.c.Metazoa and Qa.I.Ufe1) reached nine; a single divergent group (Qb.III.b.SAR) required manual adjustment and converged after 21 iterations.

#### Adaptive threshold selection using mixture modelling

To determine iteration-specific inclusion thresholds in an objective manner, the distribution of hmmsearch scores was analysed at each iteration. E-values were transformed to −log□(E-value), which compresses the wide dynamic range of HMMER E-values and renders the right-skewed score distribution approximately symmetric and amenable to Gaussian modelling, with smaller E-values (stronger hits) mapping to higher transformed values. The transformed scores were modelled using a two-component Gaussian Mixture Model (GMM), capturing the bimodal nature of the distribution: a high-scoring component corresponding to genuine subfamily homologs and a low-scoring component corresponding to non-specific hits and background noise. The relationship between score distributions, inferred thresholds and phylogenetic clustering is illustrated for the Qa.I SNAREs (Supplementary Fig. S16).

For each iteration, the Gaussian component with the highest mean score was interpreted as representing the distribution of true positive hits. The inclusion threshold was defined relative to this component as the mean minus one standard deviation, corresponding approximately to the 16th percentile of the high-scoring distribution, so that the cutoff was learned from the data at each iteration and adapted to the per-subfamily score distribution rather than relying on a fixed, arbitrary E-value threshold. An adaptive offset was applied when necessary to prevent premature convergence. Sequences exceeding this threshold were incorporated into the training set for the next iteration.

To ensure stability of the refinement process, the threshold offset was progressively adjusted when the number of newly selected sequences failed to increase between iterations.

#### Final model generation

The iterative refinement procedure was terminated when no additional sequences meeting the inclusion criteria could be identified. The final sequence set was then used to construct the definitive HMM for each SNARE group or subgroup. These optimized models were subsequently incorporated into the updated database and used for downstream annotation of SNARE domains.

### Implementation

The overall structure of the TRACEY database and web interface is summarized in Figure 1. TRACEY integrates a curated sequence collection with refined HMM-based domain models (Fig. 1A) and provides a web platform for flexible querying, visualization and data download (Fig. 1B), together with utilities for user-submitted sequence analysis and phylogeny-based exploration of SNAREs within the broader context of the vesicle-trafficking machinery in which they operate (Fig. 1C).

**Figure 1.**
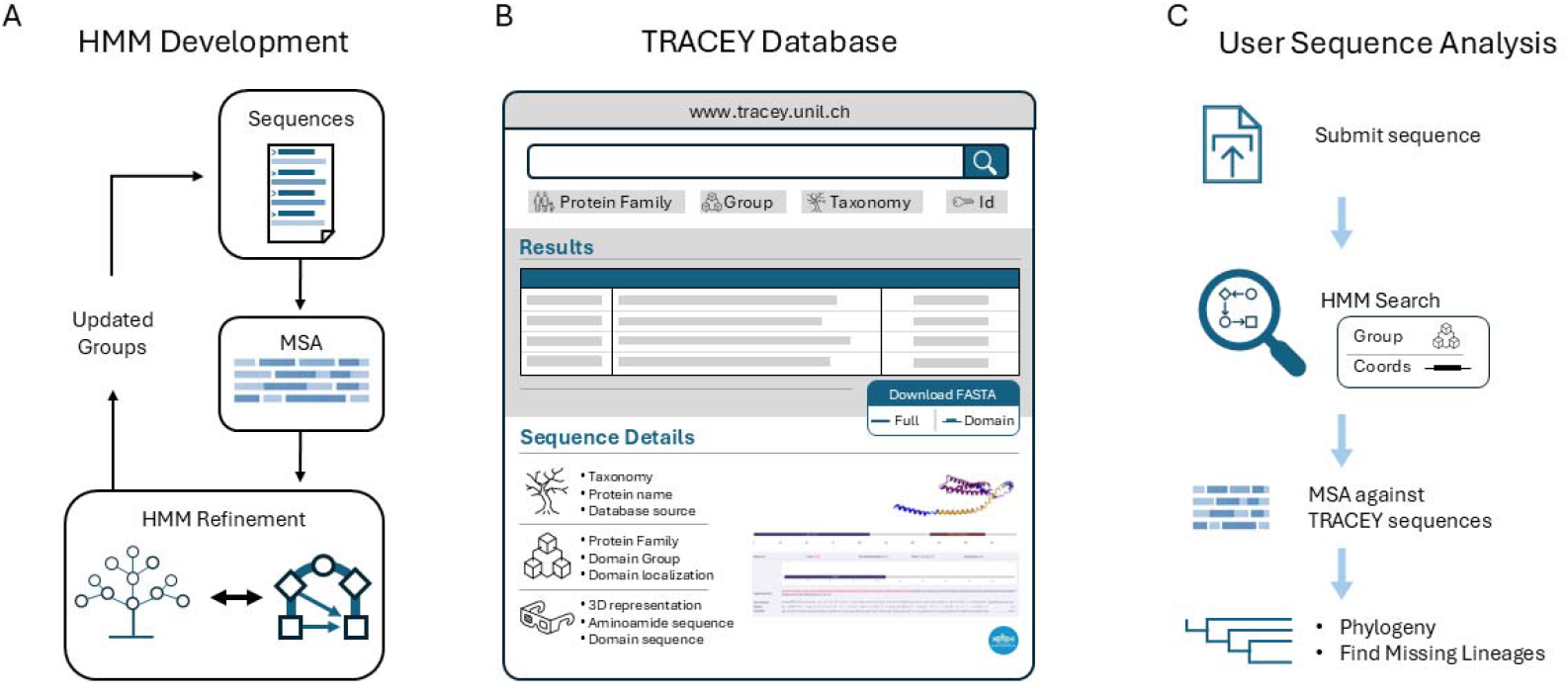
TRACEY Overview *A*. Sequence collection and HMM refinement process. *B*. TRACEY FUNCTIONALITY: Searching and filtering results, download sequences/domains in fasta format, domain details. *C*. Utilities and usability: submit new sequences, search protein domains in sequences, alignments against TRACEY sequences, phylogeny analysis, find under sampled or previously unrecognized lineages.

The new TRACEY webserver provides a flexible multi-parameter search interface that enables users to query the database by SNARE group, class or subclass, species or broader taxonomic levels, protein name, or sequence identifier. Query results are presented in a clear, structured format that facilitates rapid inspection and downstream parsing, and can be downloaded in FASTA format either as full-length protein sequences or as isolated SNARE-domain sequences. Each sequence can also be examined individually through a dedicated detail page that displays its taxonomic assignment, source database and accession, available 3D structural representation derived from the AlphaFold Protein Structure Database (Varadi et al., 2022), domain boundaries, subgroup classification, and the associated HMM-derived scores, E-values and alignment.

In addition to browsing the curated dataset, users may submit external sequences for analysis against the refined HMM library. Submitted proteins are scanned against all TRACEY HMMs, and the server returns matching profiles together with their statistical support, classification information, and predicted domain coordinates. Advanced users may request additional privileges, including the ability to generate sequence alignments against TRACEY entries or to submit novel sequences for integration into the automated HMM-based annotation workflow. User registration is used exclusively for authentication, and no personal information beyond a username, e-mail and password is stored.

#### Sequences update

All TRACEY sequences were systematically synchronized with current NCBI records to ensure consistency with publicly available sequence identifiers and versioning. For each live TRACEY entry, we first identified a similarity block consisting of all TRACEY sequences from the same species sharing ≥80% sequence identity, representing alternative versions or redundant records of the same protein.

All sequences within each similarity block were queried against NCBI using the Entrez Programming Utilities (E-utilities) to determine their current status. Based on these queries, deprecated accessions were flagged, and sequences that had been superseded by newer NCBI versions were updated accordingly. For each block, a single representative sequence corresponding to the current NCBI record was retained as the live TRACEY entry.

When a newer NCBI sequence version was identified but not yet present in TRACEY, a new database entry was generated and linked to the corresponding similarity block. This procedure ensured that TRACEY reflects the current NCBI sequence space while avoiding redundancy and preserving traceability between legacy and updated records.

## Discussion

### Comprehensive update of the TRACEY database

Synchronization of TRACEY with current NCBI sequence records resulted in extensive revisions to the underlying dataset. Of the sequences present in the original release, 3,633 accessions were identified as deprecated in NCBI and were removed from the set of live TRACEY entries, while 1,715 sequences were replaced by updated NCBI versions. In addition, 911 newly identified SNARE protein sequences absent from the original database were incorporated, reflecting the expanded availability of genomic data from previously under sampled taxa. Conversely, 1,686 sequences that had previously been flagged as non-live in TRACEY were reinstated as live entries to match their current NCBI status **(Table 1)**. Collectively, these updates substantially reshaped the database sequence collection, ensuring that TRACEY accurately reflects the current SNARE sequence space while maintaining a non-redundant and version-consistent representation of SNARE proteins across species. A complete breakdown of the sequence status transitions is provided in Supplementary Table S1.

**Table 1.**
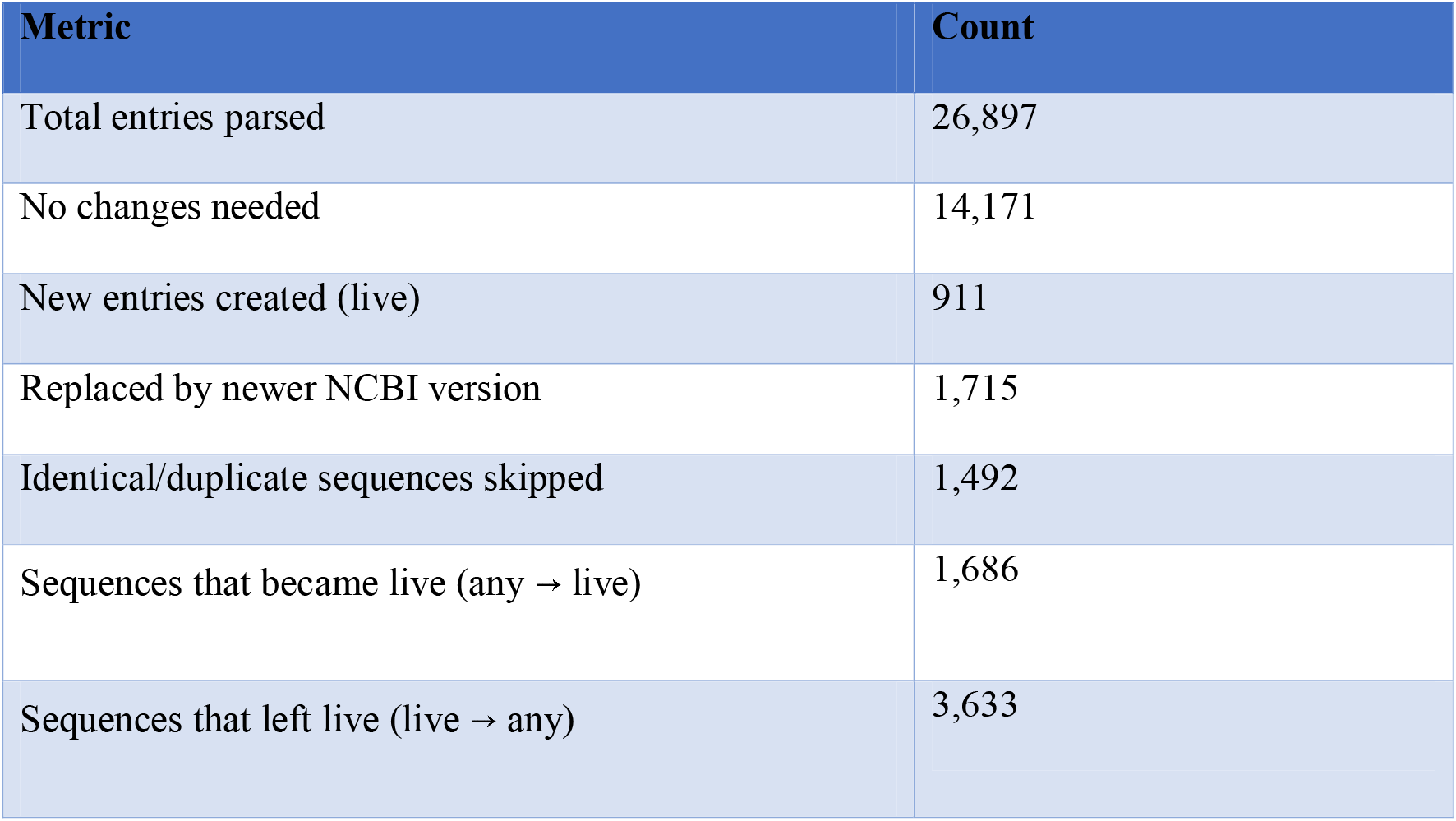
Summary of TRACEY sequence updates relative to current NCBI status.

### Improved SNARE domain detection using optimized HMM profiles

The refinement of the Hidden Markov Models (HMMs) has led to significant improvements in their performance compared to previous models. Using an enhanced methodology for HMM construction, we successfully developed 83 HMMs targeting SNARE groups and subgroups from which 43 were not defined within the older set of HMMs. This advancement allowed for a more precise categorization of sequences within these previously uncharacterized subgroups. Furthermore, a comparative analysis of the E-values obtained for sequences present in both the old and new datasets, in which the legacy and optimized HMMs were scanned against the same sequence set so that E-values are directly comparable, demonstrates the superiority of the updated models (Fig. 2A); at least ∼75% of sequences in every overlapping group exhibited improved scoring with the new HMMs, with the R.I group showing the lowest improvement rate (Fig. 2B). This finding not only underscores the efficacy of our refined approach but also highlights the potential for more accurate predictions in the classification of SNARE proteins. Final HMM-based classifications across all SNARE groups are visualized on domain-based phylogenetic trees (Supplementary Figs. S1–S15).

**Figure 2.**
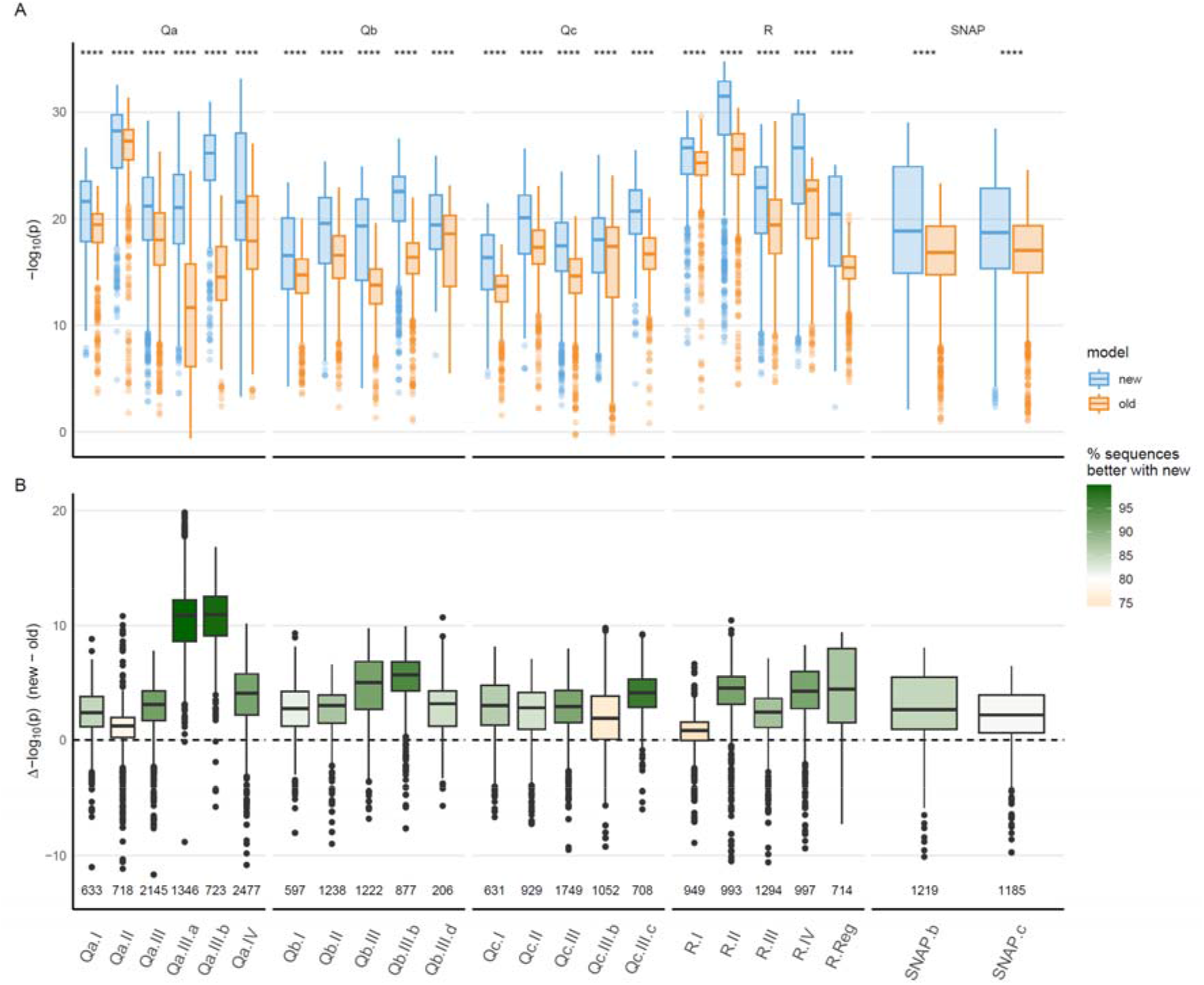
Performance comparison of optimized and legacy HMMs for SNARE domain detection. Comparison between newly optimized and previous sets of Hidden Markov Models (HMMs) for SNARE domain detection across groups shared between both model sets. For every group, the new and legacy HMMs were scanned against the same sequence set, so that the search space (and hence the E-value normalization) was identical for both models; the paired per-sequence comparison is therefore unaffected by database size. *A*. Distribution of E-values obtained for each group using the new (blue) and previous (orange) HMMs. Boxplots summarize score distributions across sequences. Statistical significance between model performances was assessed using a Wilcoxon rank-sum test, with significance levels indicated above each comparison. *B*. ΔE-value distribution calculated as the difference between logarithmic E-values obtained with the new and old HMMs for the same sequence (Δ = new − old). Positive Δ values indicate improved domain recognition by the optimized HMM. Bar coloration reflects the proportion of sequences within each group achieving better scores with the new model. Total number of sequences per group is indicated below each box. The R.I group showed the lowest improvement rate, with approximately 75% of sequences scoring better under the updated HMM.

### Limitations

Several aspects of this release should be interpreted with care. First, the comparison between optimized and legacy HMMs was performed on the curated TRACEY domain set, which also informed model construction; held-out evaluation on sequences excluded from training would provide a more stringent estimate of generalization. Second, the iterative procedure was tuned to limit low-confidence recruitment, but specificity was not benchmarked against an explicit set of non-SNARE decoy sequences. Third, because the collection is synchronized to current NCBI records, coverage remains bounded by the taxa represented there and will require periodic resynchronization. We regard these as directions for further validation rather than limitations on the curated content released here.

### Future development of the TRACEY resource

Future developments of TRACEY will focus on two main areas. First, the database will be expanded to include sequences from currently missing or underrepresented lineages, such as members of the SAR supergroup (Stramenopiles, Alveolata and Rhizaria), as well as other under-represented eukaryotic lineages such as Metamonada and early-diverging fungi. The sparse representation of these lineages is already apparent in the refinement procedure: the only group requiring extended, manually adjusted iteration (Qb.III.b.SAR) is itself a SAR subgroup, where the limited number of available sequences yields a flatter score distribution and slow per-iteration recruitment. Expanding SAR coverage is therefore expected both to improve annotation of these lineages and to stabilize model refinement for the corresponding subgroups. Second, the HMM curation pipeline developed in this work will be extended to other protein families beyond SNAREs. Together, the expanded sequence collection, the refined HMM library and the redesigned interface make TRACEY a current, reproducible and broadly accessible reference for the annotation and comparative analysis of SNARE protein domains.

## Acknowledgements

We acknowledge the support of the Centre informatique (Ci) of the University of Lausanne for providing server infrastructure and technical support for hosting the TRACEY database. We also thank Tobias Klöpper, who developed the original database architecture that formed the foundation of the present TRACEY resource, and Nickias Kienle for creating the initial TRACEY web interface and implementing early versions of the data analysis code.

## Author contributions

Carlos Pulido-Quetglas: Conceptualization, Data curation, Formal analysis, Methodology, Software, Validation, Visualization, Writing, original draft. Dirk Fasshauer: Conceptualization, Supervision, Writing, review & editing

## Supplementary data

Supplementary Data are available at Bioinformatics online.

## Data availability

TRACEY is publicly available at https://tracey.unil.ch. Sequence data underlying TRACEY are derived from public repositories, primarily NCBI, and accession identifiers are provided within the database. The optimized HMM models are accessible through the TRACEY web interface and can be used to analyse user-submitted sequences, but are available for download only upon request. Phylogenetic trees and classification results generated in this study are available as Supplementary Data published alongside this article.

