## Supplementary material for "TRACEY: an updated resource for SNARE protein domain annotation with improved HMMs and expanded sequence coverage"

**Supplementary Data – Index**

- **Supplementary Figure S1.** Domain‑based phylogenetic tree of Qa.I SNAREs highlighting HMM‑based subgroup classifications.
- **Supplementary Figure S2.** Domain‑based phylogenetic tree of Qa.II SNAREs highlighting HMM‑based subgroup classifications.
- **Supplementary Figure S3.** Domain‑based phylogenetic tree of Qa.III SNAREs highlighting HMM‑based subgroup classifications.
- **Supplementary Figure S4.** Domain‑based phylogenetic tree of Qa.IV SNAREs highlighting HMM‑based subgroup classifications.
- **Supplementary Figure S5.** Domain‑based phylogenetic tree of Qb.I SNAREs highlighting HMM‑based subgroup classifications.
- **Supplementary Figure S6.** Domain‑based phylogenetic tree of Qb.II SNAREs highlighting HMM‑based subgroup classifications.
- **Supplementary Figure S7.** Domain‑based phylogenetic tree of Qb.III SNAREs highlighting HMM‑based subgroup classifications.
- **Supplementary Figure S8.** Domain‑based phylogenetic tree of Qc.I SNAREs highlighting HMM‑based subgroup classifications.
- **Supplementary Figure S9.** Domain‑based phylogenetic tree of Qc.II SNAREs highlighting HMM‑based subgroup classifications.
- **Supplementary Figure S10.** Domain‑based phylogenetic tree of Qc.III SNAREs highlighting HMM‑based subgroup classifications.
- **Supplementary Figure S11.** Domain‑based phylogenetic tree of SNAP SNAREs highlighting HMM‑based subgroup classifications.
- **Supplementary Figure S12.** Domain‑based phylogenetic tree of R.I SNAREs highlighting HMM‑based subgroup classifications.
- **Supplementary Figure S13.** Domain‑based phylogenetic tree of R.III SNAREs highlighting HMM‑based subgroup classifications.
- **Supplementary Figure S14.** Domain‑based phylogenetic tree of R.IV SNAREs highlighting HMM‑based subgroup classifications.
- **Supplementary Figure S15.** Domain‑based phylogenetic tree of R.Reg SNAREs highlighting HMM‑based subgroup classifications.
- **Supplementary Figure S16.** Definition of subgroup boundaries during HMM optimization for Qa.I SNAREs using E‑value distributions and score‑mapped phylogenetic trees.
- **Supplementary Table 1.** Detailed breakdown of sequence status transitions in TRACEY.

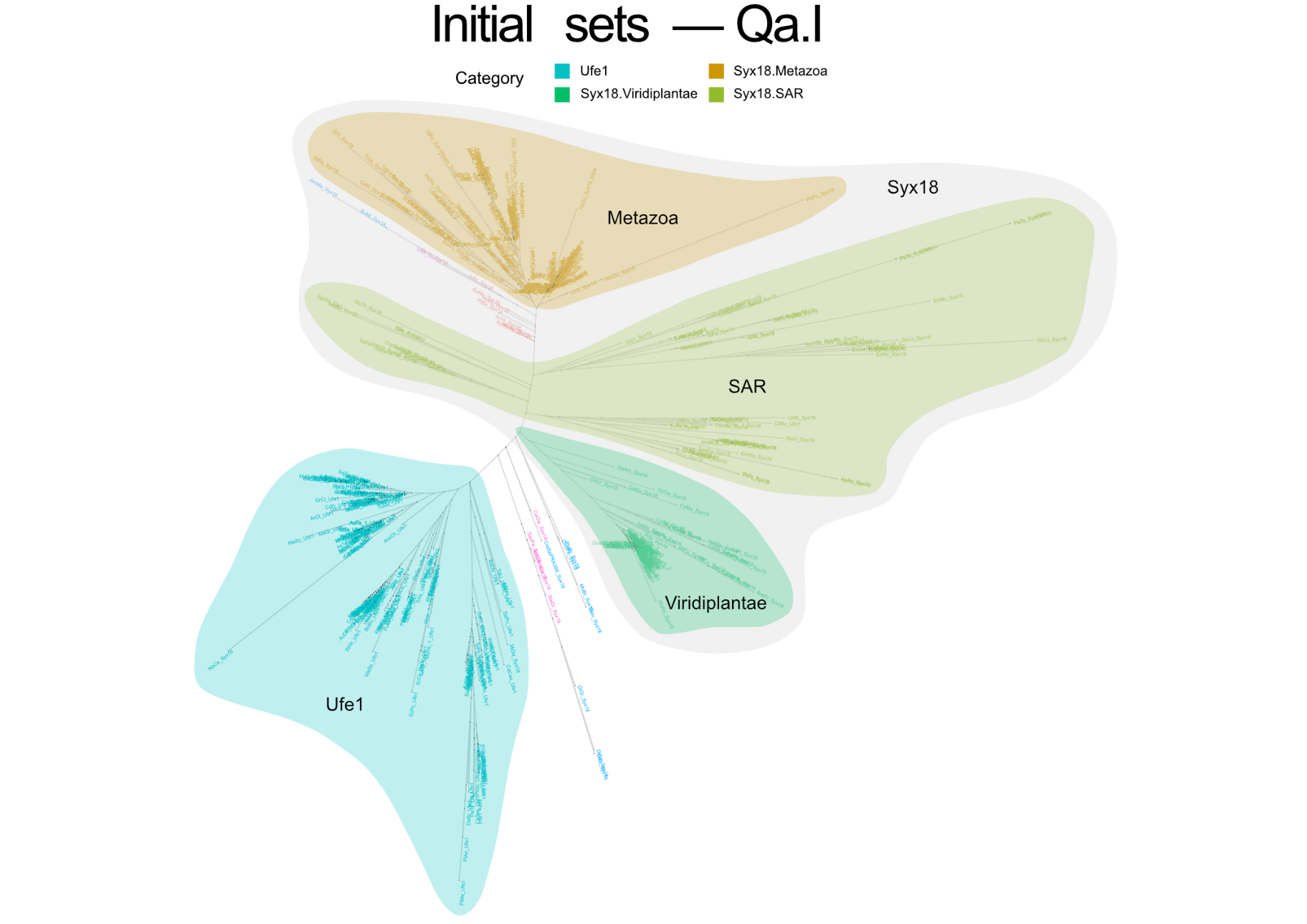

**Supplementary Figure S1. HMM‑based classification of Qa.I SNAREs.** Domain‑based phylogenetic tree of Qa.I SNARE sequences, with branches coloured according to subgroup assignments obtained using the optimized HMM profiles. Highlighted clades correspond to lineage‑ and subclass‑specific groups defined during HMM refinement, illustrating the consistency between phylogenetic structure and HMM‑based classification.

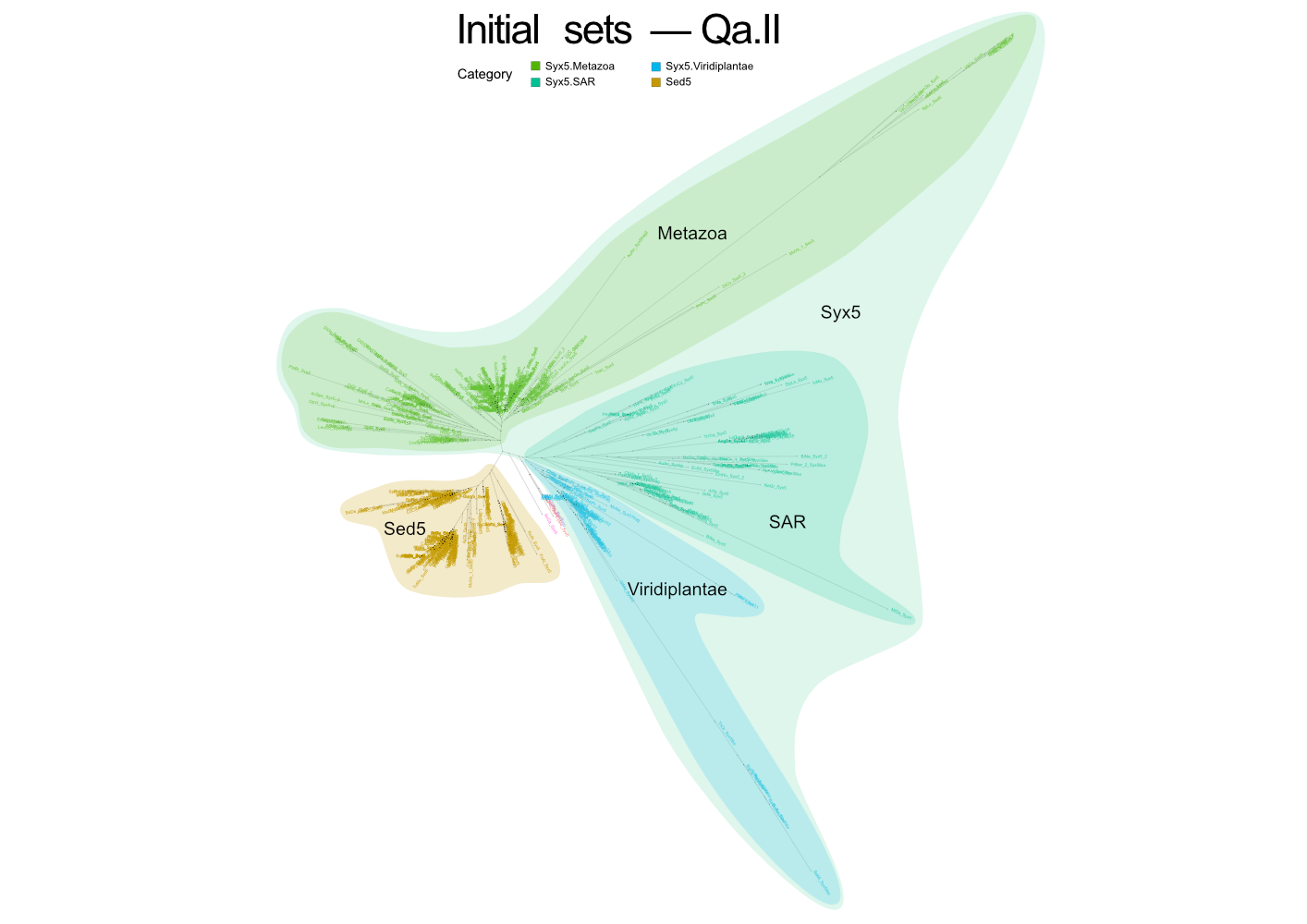

**Supplementary Figure S2. HMM‑based classification of Qa.II SNAREs.** Domain‑based phylogenetic tree of Qa.II SNARE sequences, with branches coloured according to subgroup assignments obtained using the optimized HMM profiles. Highlighted clades correspond to lineage‑ and subclass‑specific groups defined during HMM refinement, illustrating the consistency between phylogenetic structure and HMM‑based classification.

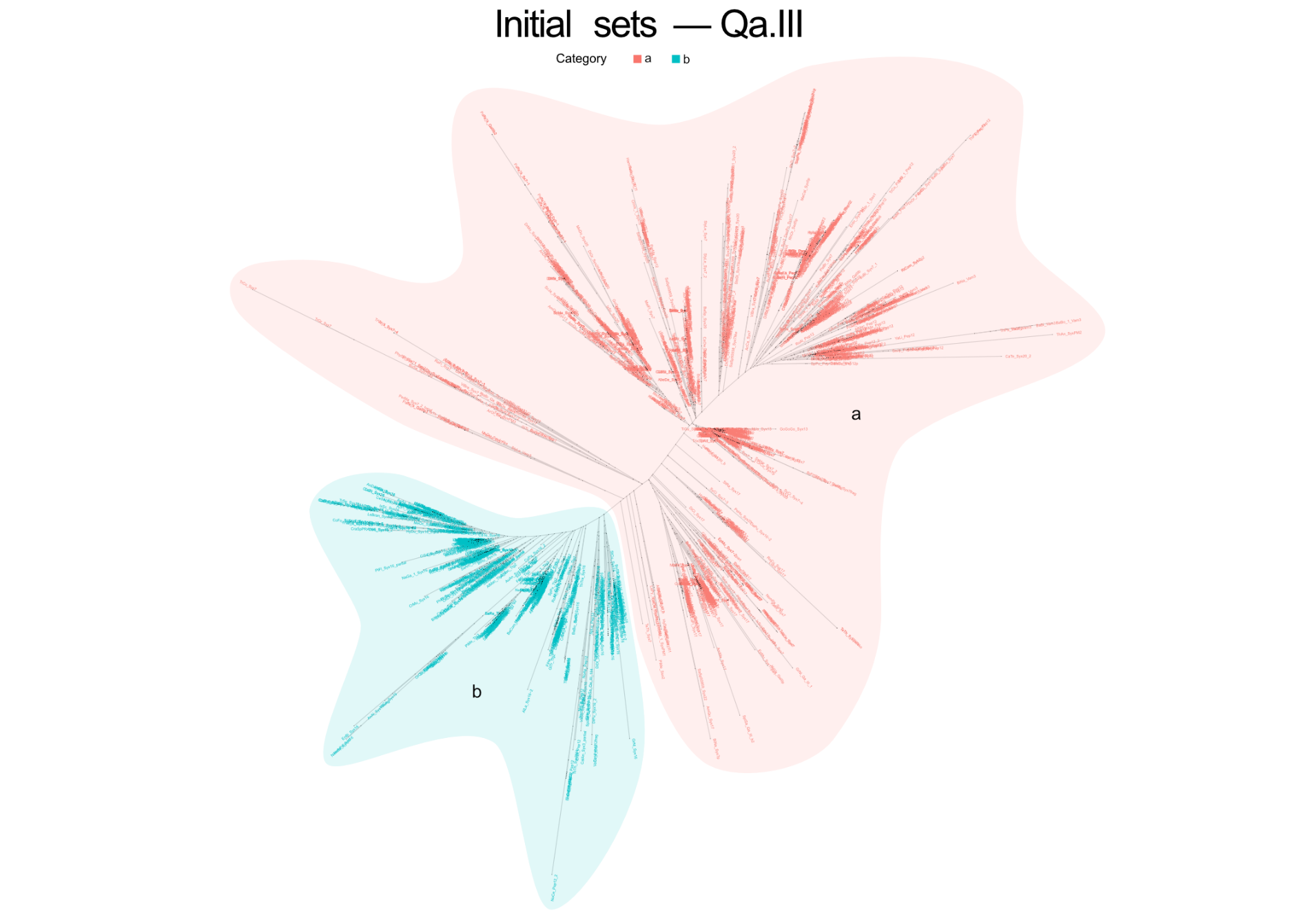

**Supplementary Figure S3. HMM‑based classification of Qa.III SNAREs.** Domain‑based phylogenetic tree of Qa.III SNARE sequences, with branches coloured according to subgroup assignments obtained using the optimized HMM profiles. Highlighted clades correspond to lineage‑ and subclass‑specific groups defined during HMM refinement, illustrating the consistency between phylogenetic structure and HMM‑based classification.

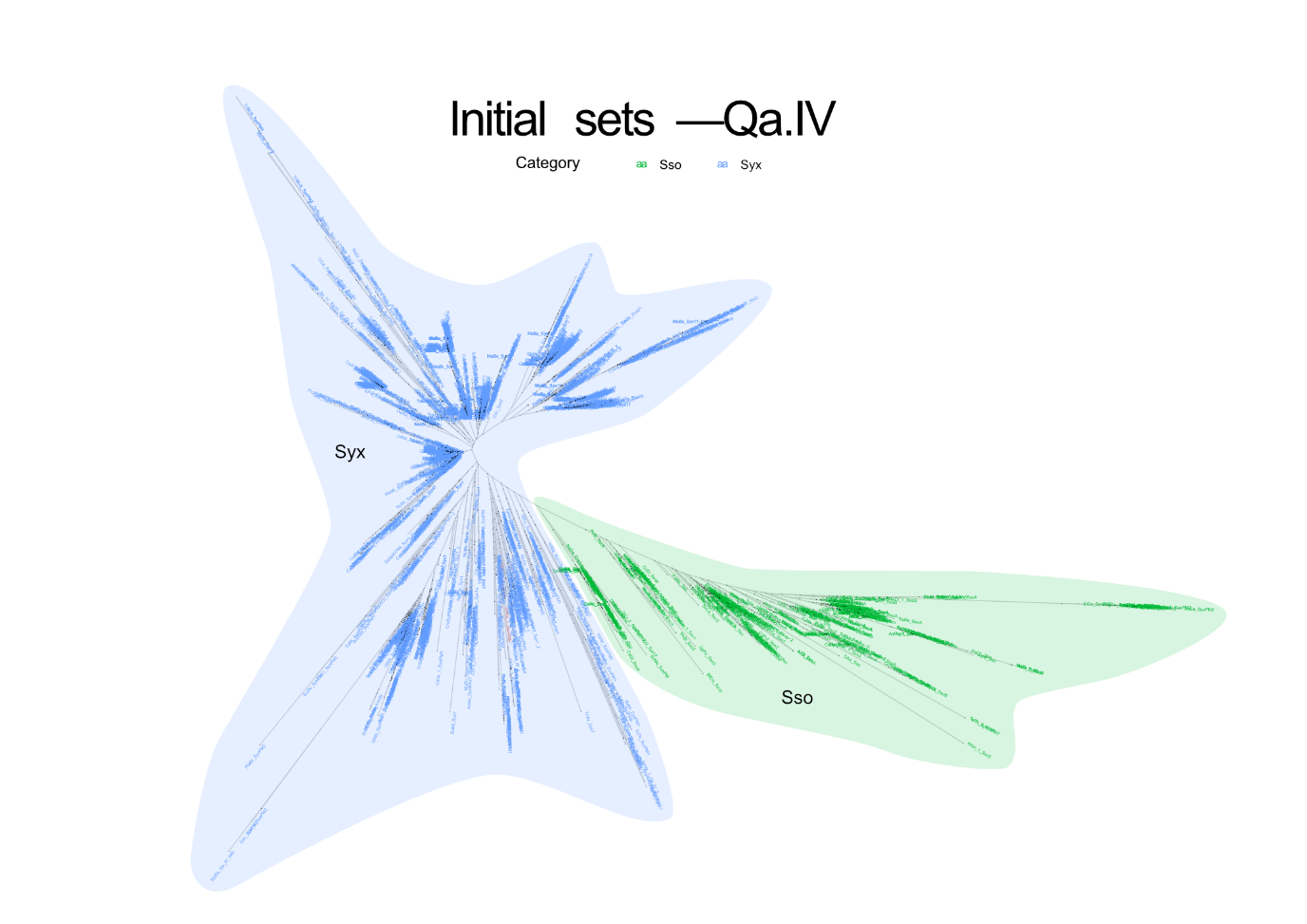

**Supplementary Figure S4. HMM‑based classification of Qa.IV SNAREs.** Domain‑based phylogenetic tree of Qa.IV SNARE sequences, with branches coloured according to subgroup assignments obtained using the optimized HMM profiles. Highlighted clades correspond to lineage‑ and subclass‑specific groups defined during HMM refinement, illustrating the consistency between phylogenetic structure and HMM‑based classification.

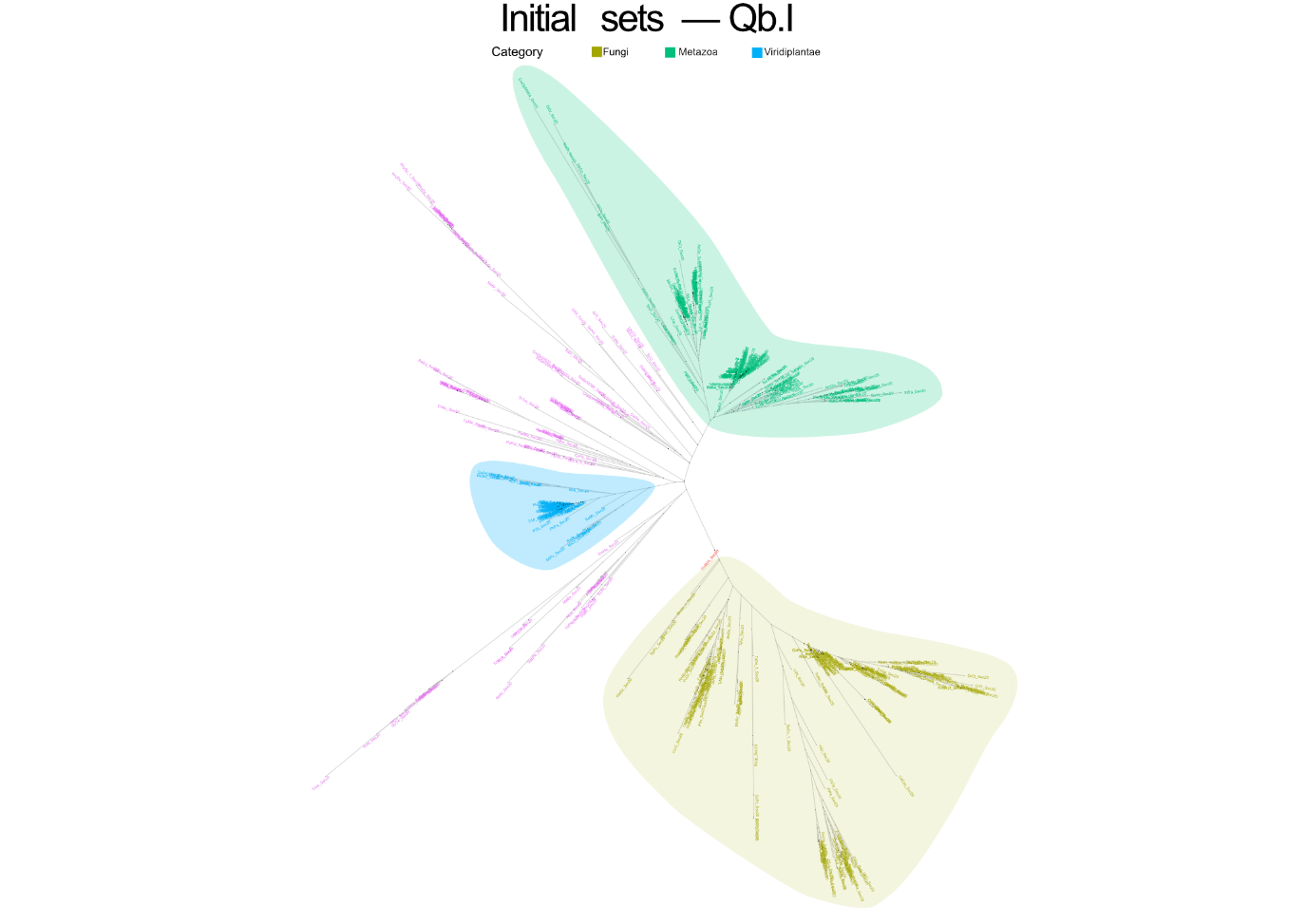

**Supplementary Figure S5. HMM‑based classification of Qb.I SNAREs.** Domain‑based phylogenetic tree of Qb.I SNARE sequences, with branches coloured according to subgroup assignments obtained using the optimized HMM profiles. Highlighted clades correspond to lineage‑ and subclass‑specific groups defined during HMM refinement, illustrating the consistency between phylogenetic structure and HMM‑based classification.

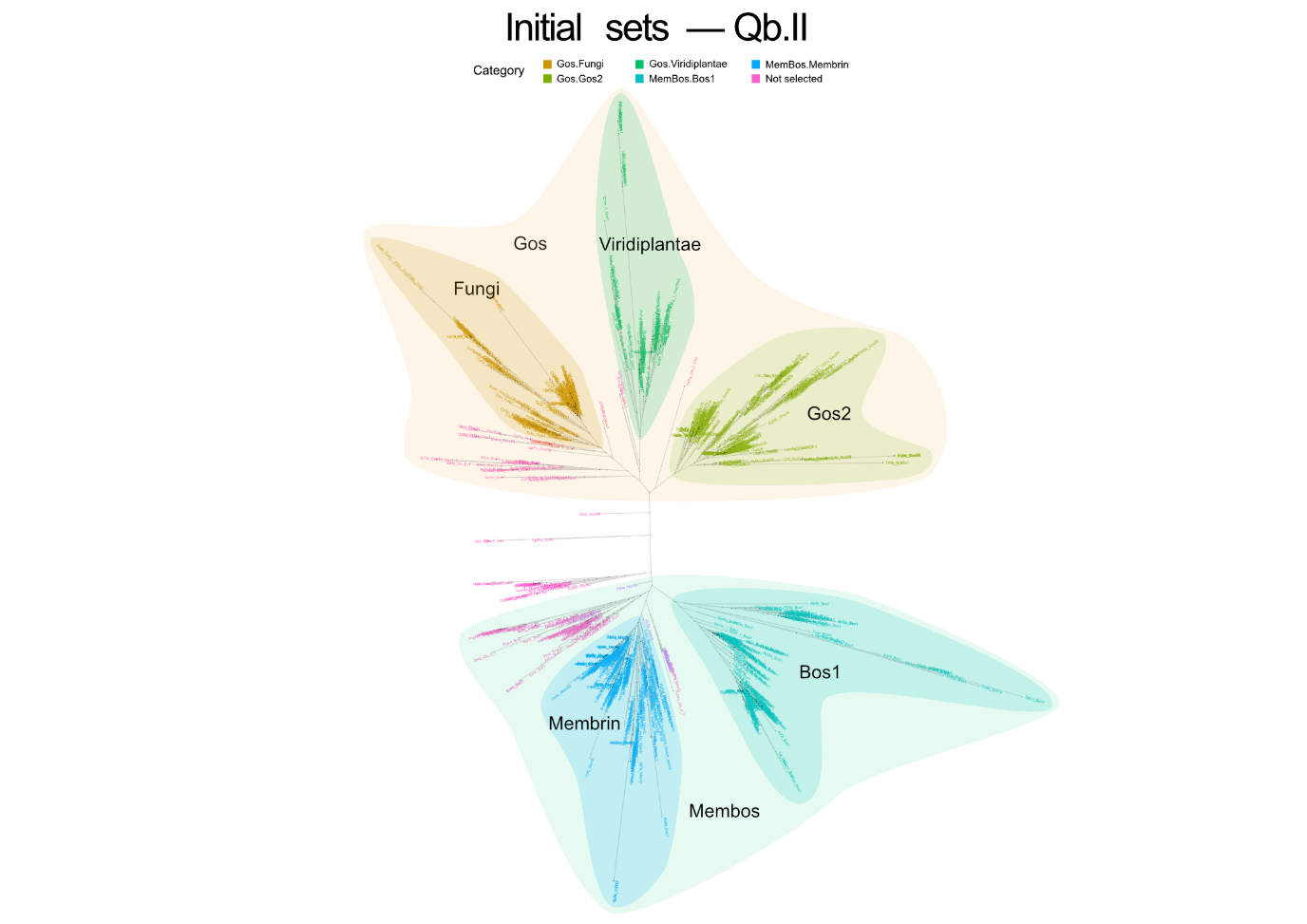

**Supplementary Figure S6. HMM‑based classification of Qb.II SNAREs.** Domain‑based phylogenetic tree of Qb.II SNARE sequences, with branches coloured according to subgroup assignments obtained using the optimized HMM profiles. Highlighted clades correspond to lineage‑ and subclass‑specific groups defined during HMM refinement, illustrating the consistency between phylogenetic structure and HMM‑based classification.

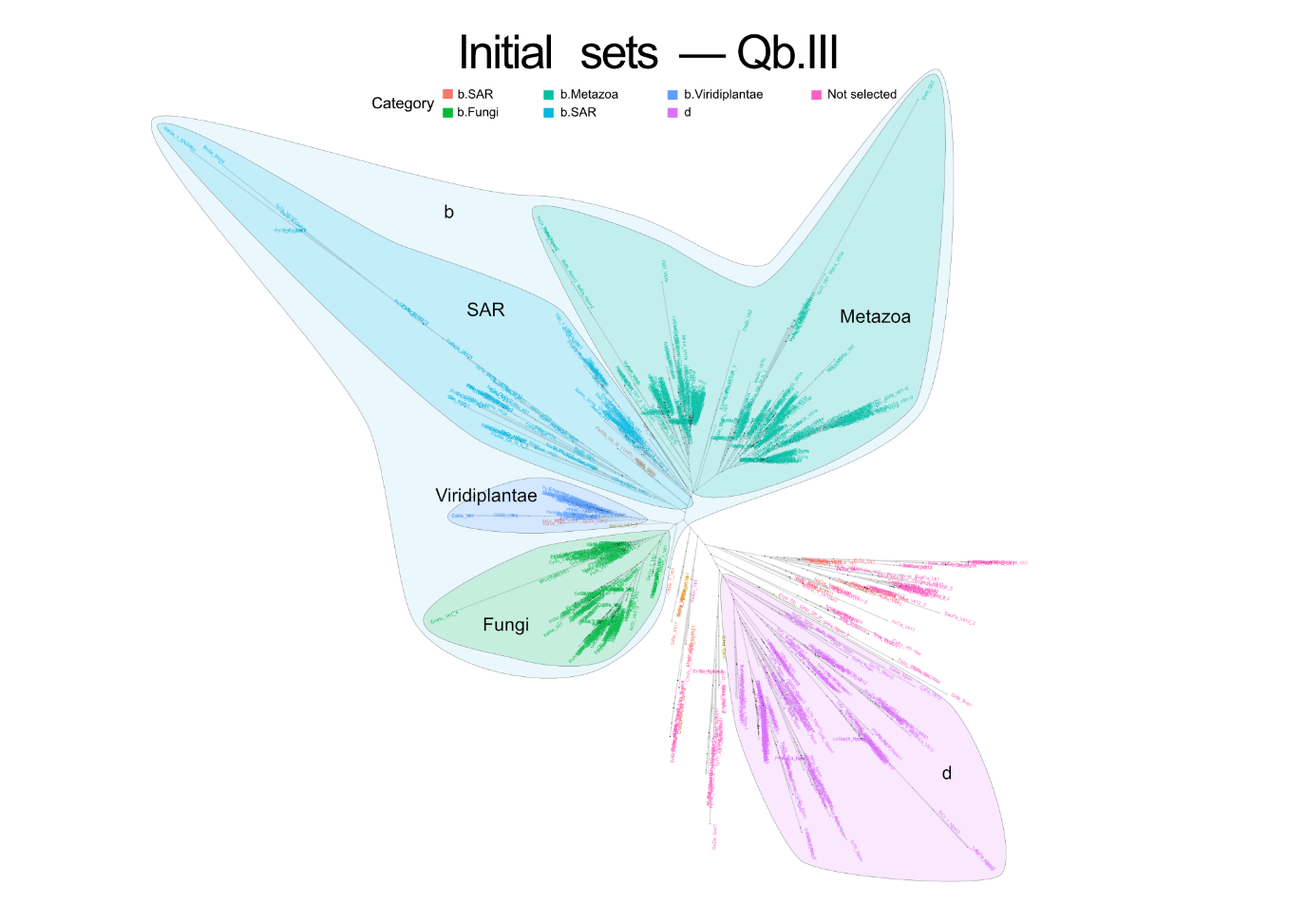

**Supplementary Figure S7. HMM‑based classification of Qb.III SNAREs.** Domain‑based phylogenetic tree of Qb.III SNARE sequences, with branches coloured according to subgroup assignments obtained using the optimized HMM profiles. Highlighted clades correspond to lineage‑ and subclass‑specific groups defined during HMM refinement, illustrating the consistency between phylogenetic structure and HMM‑based classification.

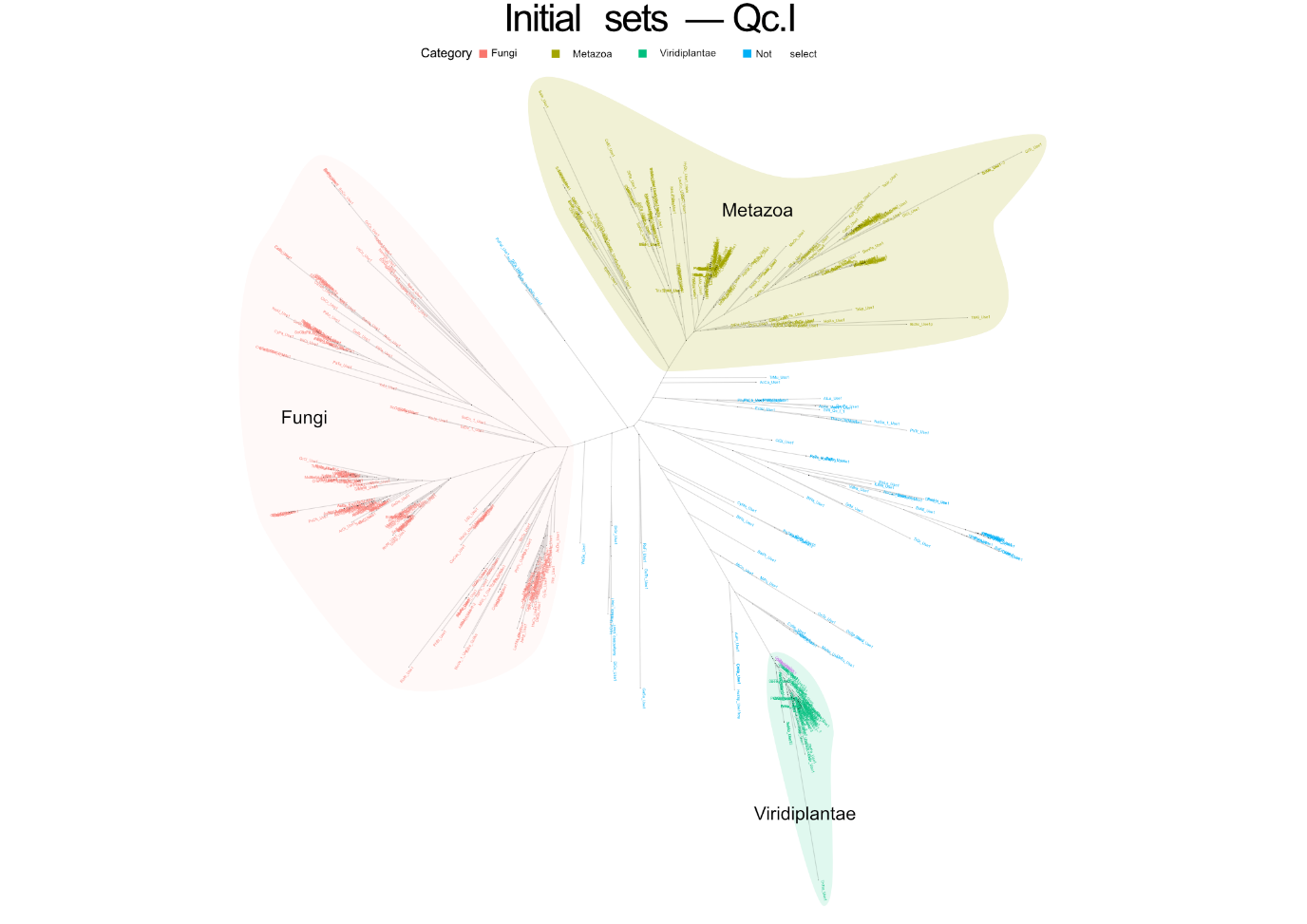

**Supplementary Figure S8. HMM‑based classification of Qc.I SNAREs.** Domain‑based phylogenetic tree of Qc.I SNARE sequences, with branches coloured according to subgroup assignments obtained using the optimized HMM profiles. Highlighted clades correspond to lineage‑ and subclass‑specific groups defined during HMM refinement, illustrating the consistency between phylogenetic structure and HMM‑based classification.

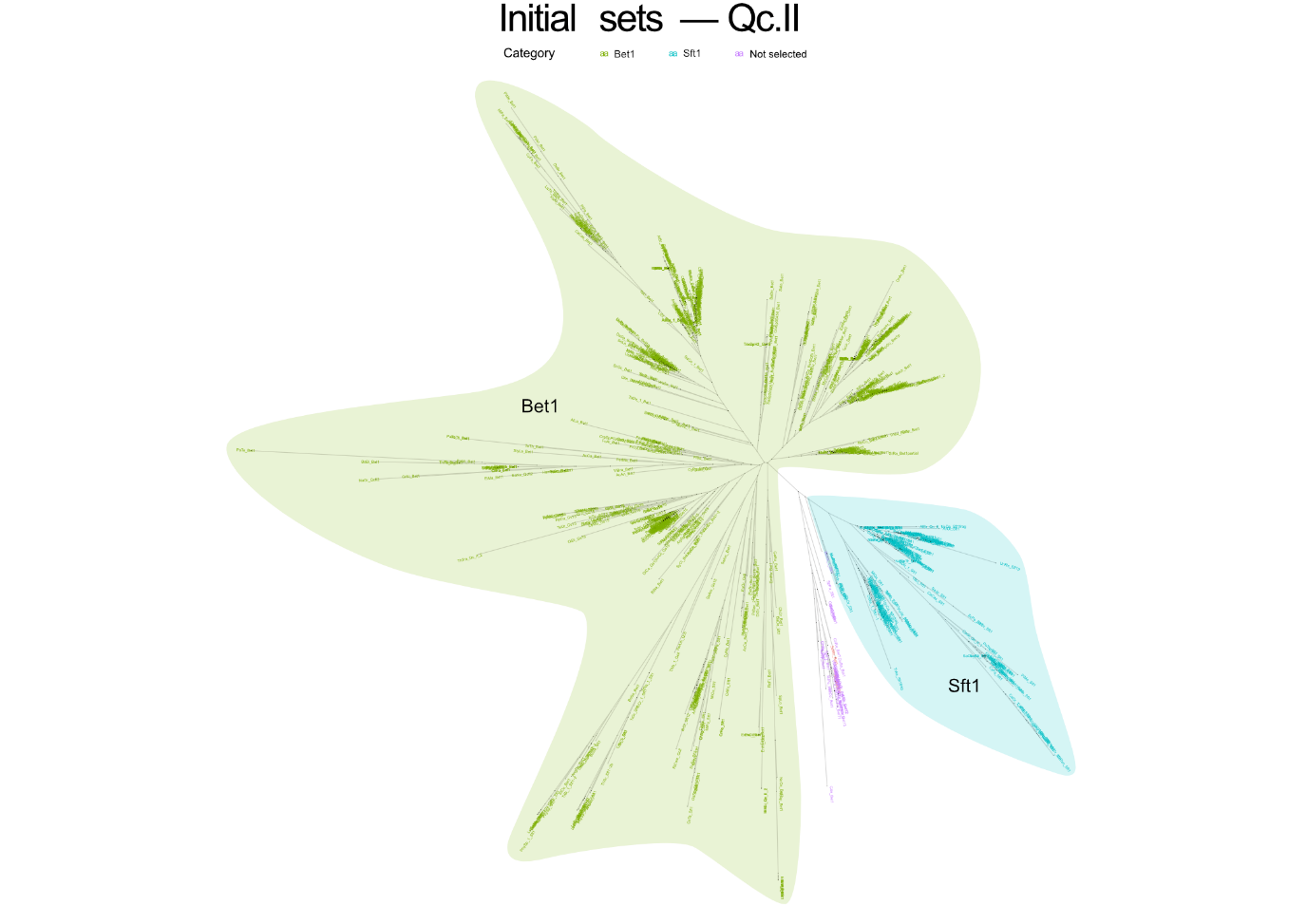

**Supplementary Figure S9. HMM‑based classification of Qc.II SNAREs.** Domain‑based phylogenetic tree of Qc.II SNARE sequences, with branches coloured according to subgroup assignments obtained using the optimized HMM profiles. Highlighted clades correspond to lineage‑ and subclass‑specific groups defined during HMM refinement, illustrating the consistency between phylogenetic structure and HMM‑based classification.

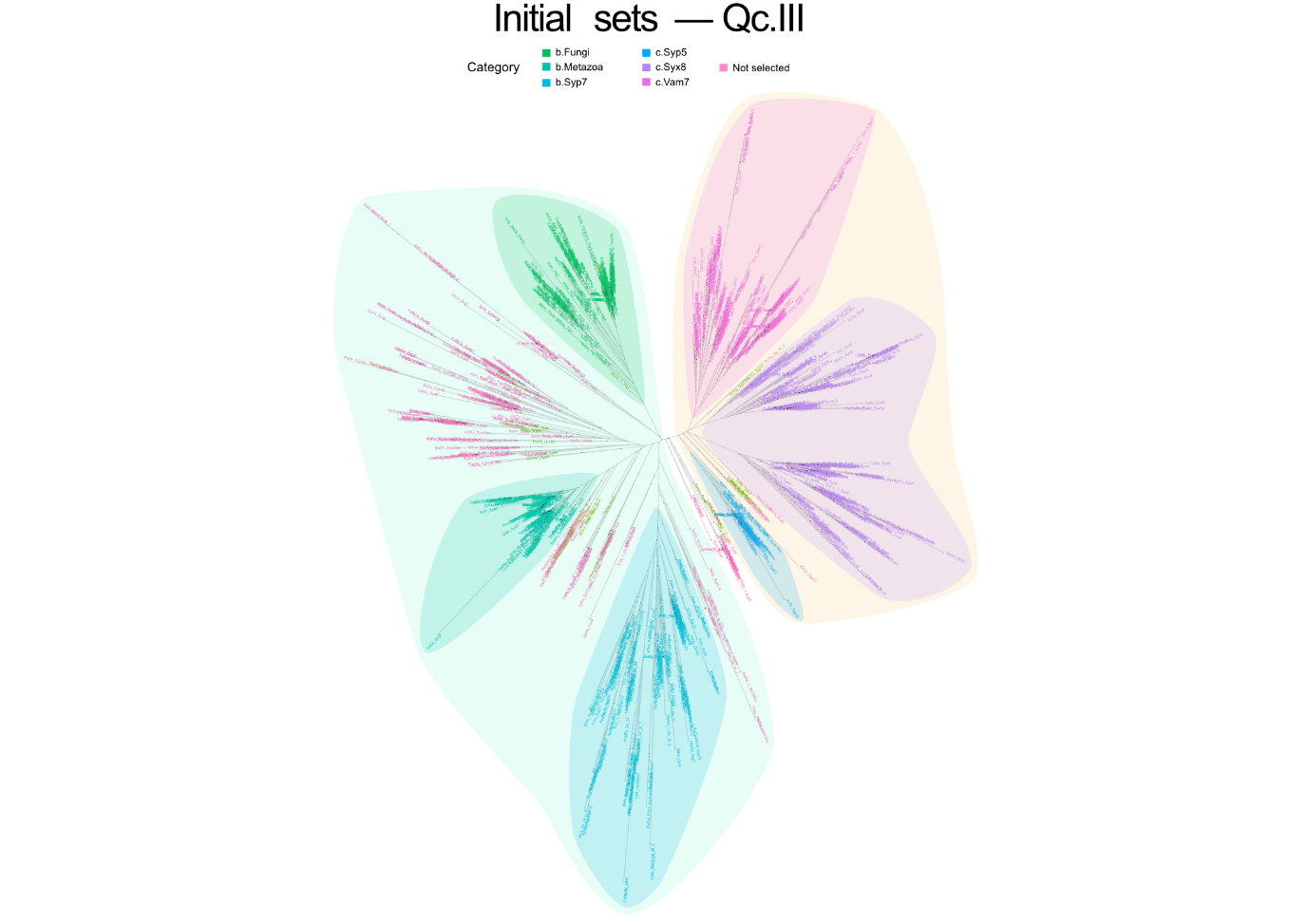

**Supplementary Figure S10. HMM‑based classification of Qc.III SNAREs.** Domain‑based phylogenetic tree of Qc.III SNARE sequences, with branches coloured according to subgroup assignments obtained using the optimized HMM profiles. Highlighted clades correspond to lineage‑ and subclass‑specific groups defined during HMM refinement, illustrating the consistency between phylogenetic structure and HMM‑based classification.

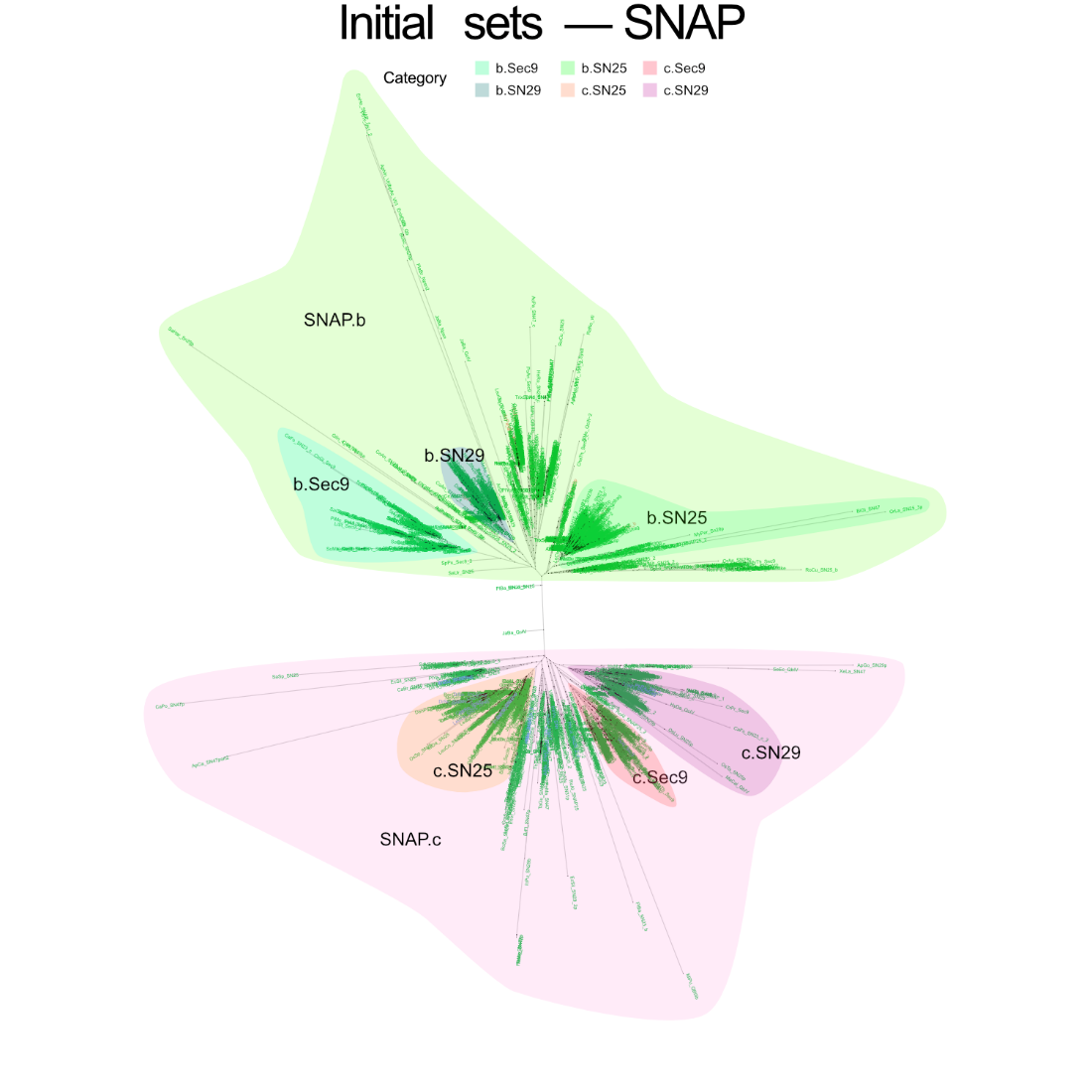

**Supplementary Figure S11. HMM‑based classification of SNAP SNAREs.** Domain‑based phylogenetic tree of SNAP SNARE sequences, with branches coloured according to subgroup assignments obtained using the optimized HMM profiles. Highlighted clades correspond to lineage‑ and subclass‑specific groups defined during HMM refinement, illustrating the consistency between phylogenetic structure and HMM‑based classification.

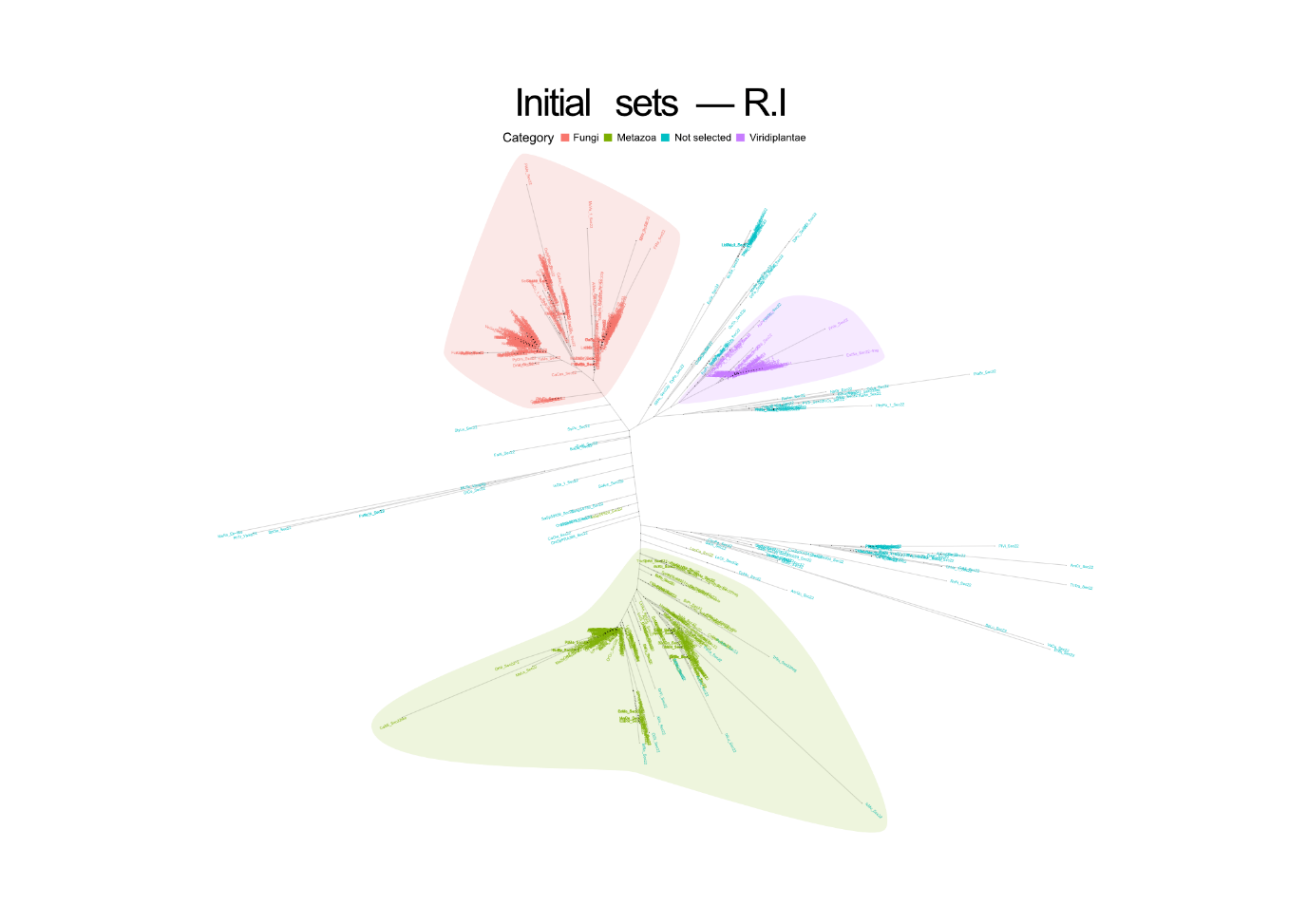

**Supplementary Figure S12. HMM‑based classification of R.I SNAREs.** Domain‑based phylogenetic tree of R.I SNARE sequences, with branches coloured according to subgroup assignments obtained using the optimized HMM profiles. Highlighted clades correspond to lineage‑ and subclass‑specific groups defined during HMM refinement, illustrating the consistency between phylogenetic structure and HMM‑based classification.

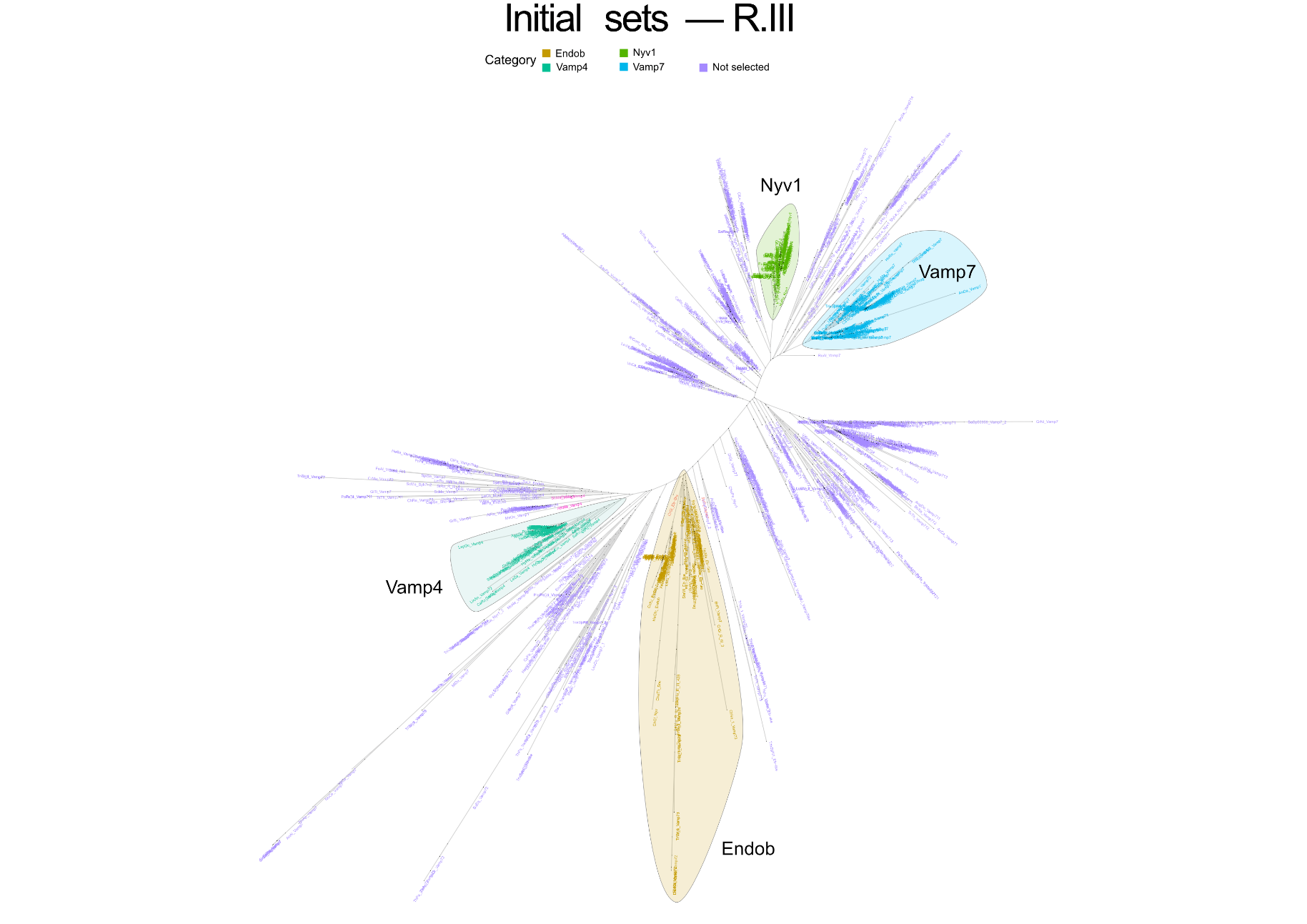

**Supplementary Figure S13. HMM‑based classification of R.III SNAREs.** Domain‑based phylogenetic tree of R.III SNARE sequences, with branches coloured according to subgroup assignments obtained using the optimized HMM profiles. Highlighted clades correspond to lineage‑ and subclass‑specific groups defined during HMM refinement, illustrating the consistency between phylogenetic structure and HMM‑based classification.

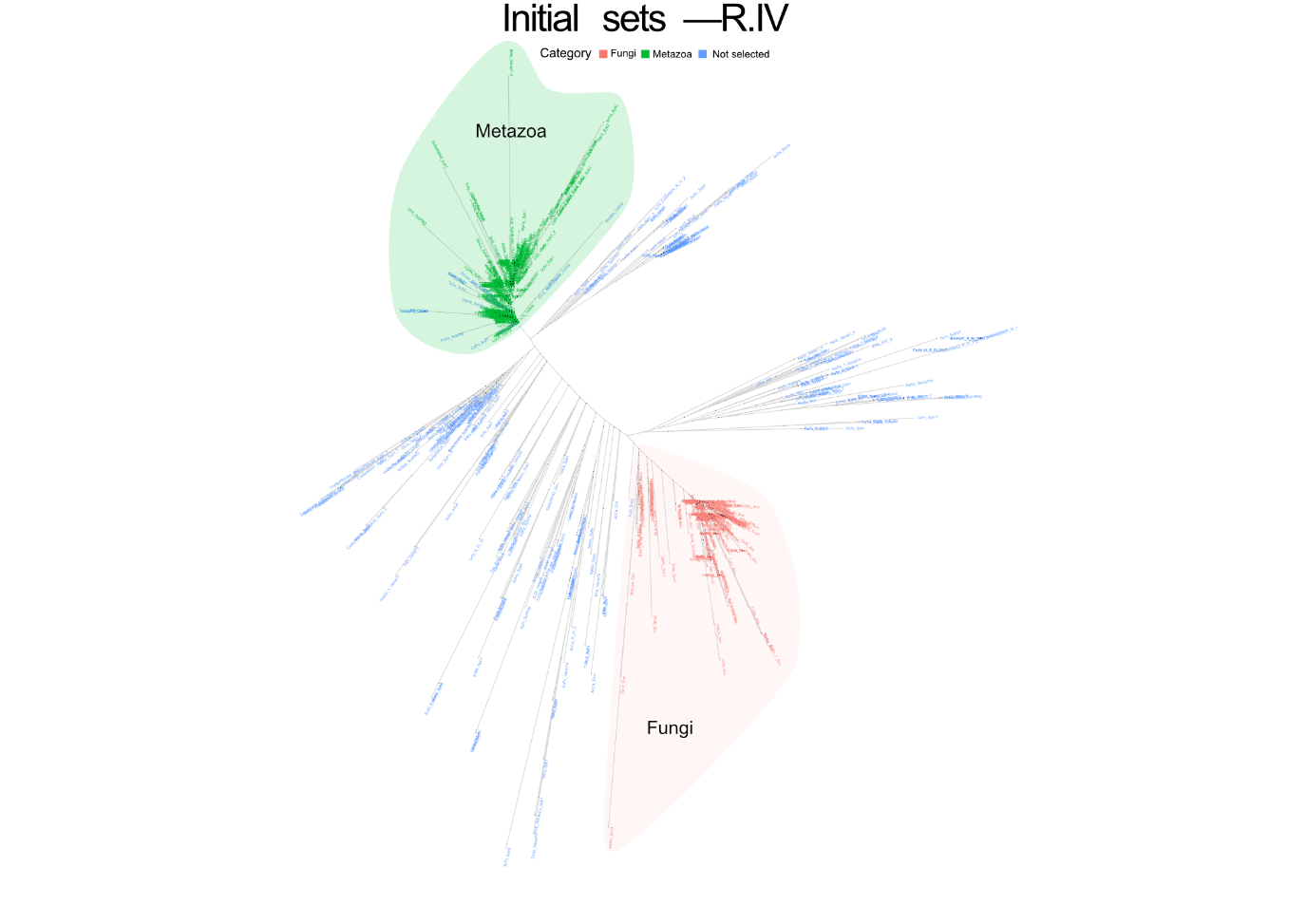

**Supplementary Figure S14. HMM‑based classification of R.IV SNAREs.** Domain‑based phylogenetic tree of R.IV SNARE sequences, with branches coloured according to subgroup assignments obtained using the optimized HMM profiles. Highlighted clades correspond to lineage‑ and subclass‑specific groups defined during HMM refinement, illustrating the consistency between phylogenetic structure and HMM‑based classification.

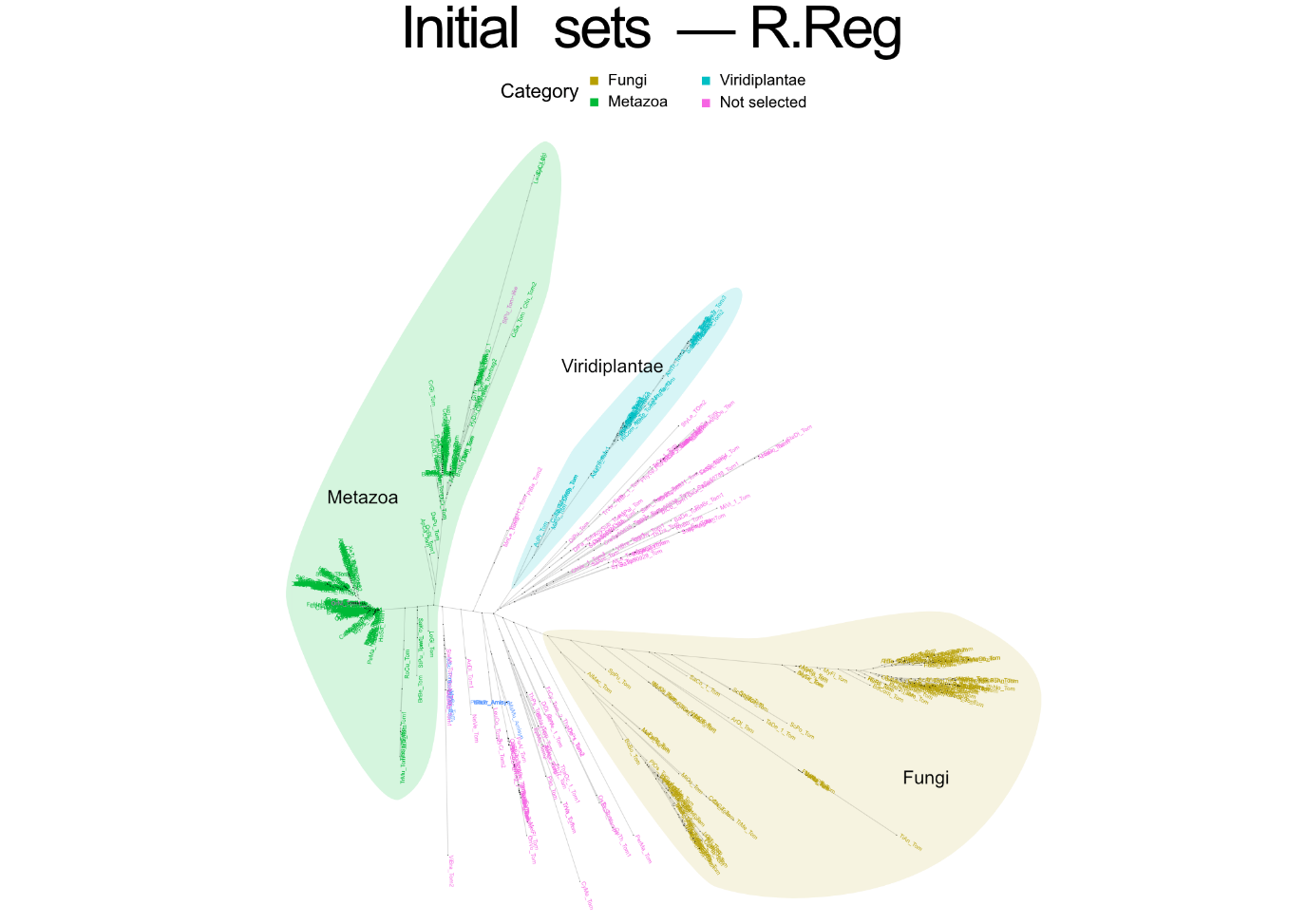

**Supplementary Figure S15. HMM‑based classification of R.Reg SNAREs.** Domain‑based phylogenetic tree of R.Reg SNARE sequences, with branches coloured according to subgroup assignments obtained using the optimized HMM profiles. Highlighted clades correspond to lineage‑ and subclass‑specific groups defined during HMM refinement, illustrating the consistency between phylogenetic structure and HMM‑based classification.

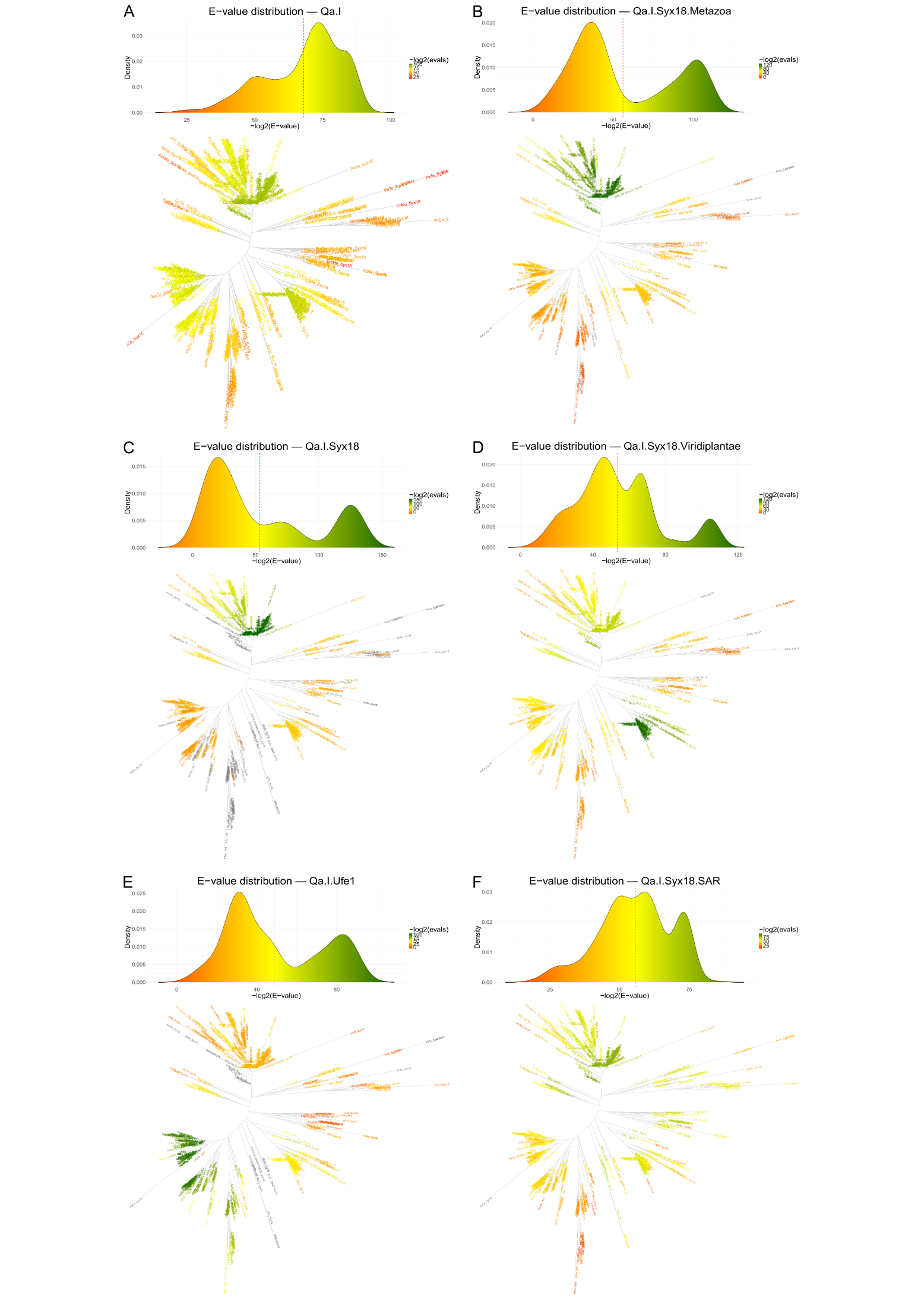

**Supplementary Figure S16. Definition of subgroup boundaries during HMM optimization for Qa.I SNAREs.** Six panels illustrate the relationship between HMM E‑value distributions and phylogenetic clustering within the Qa.I SNARE group. For each subgroup, score distributions obtained during iterative HMM refinement are shown alongside domain‑based phylogenetic trees, with sequences coloured according to −log₂(E‑value) scores. These visualizations document the criteria used to define inclusion thresholds and subgroup boundaries and illustrate the correspondence between statistical support and phylogenetic coherence during model optimization.

**Table S1.** Detailed breakdown of sequence status transitions (8,685 total) identified during synchronization with current NCBI status.

| Status transition | Count |
| --- | --- |
| live → suppressed | 2,379 |
| ignore → suppressed | 2,308 |
| ignore → live | 1,651 |
| live → replaced | 1,059 |
| ignore → replaced | 519 |
| live → dead | 194 |
| ignore → dead | 186 |
| suppressed → ignore | 168 |
| unknown → ignore | 67 |
| unknown → suppressed | 35 |
| unknown → replaced | 34 |
| suppressed → replaced | 23 |
| suppressed → live | 17 |
| unknown → live | 17 |
| suppressed → dead | 16 |
| unknown → dead | 5 |
| dead → suppressed | 2 |
| live → withdrawn | 1 |
| dead → ignore | 1 |
| replaced → dead | 1 |
| dead → replaced | 1 |
| dead → live | 1 |
| Total | 8,685 |

**Supplementary Table S2. Taxonomic coverage of the TRACEY dataset by kingdom and eukaryotic supergroup.**

| Taxonomic group | Species | Sequences |
| --- | --- | --- |
| Metazoa | 498 | 8,285 |
| Fungi | 392 | 5,511 |
| Viridiplantae | 108 | 1,635 |
| SAR | 83 | 1,469 |
| Discoba | 31 | 553 |
| Opisthokonta (protist) | 19 | 398 |
| Amoebozoa | 18 | 358 |
| Metamonada | 11 | 181 |
| Archaeplastida (red algae/glaucophytes) | 7 | 79 |
| Haptista | 4 | 42 |
| Malawimonada | 2 | 20 |
| Cryptista | 1 | 27 |
| Unclassified | 14 | 357 |
| Total | 1,188 | 18,915 |
